# Flipped Over U: Structural Basis for dsRNA Cleavage by the SARS-CoV-2 Endoribonuclease

**DOI:** 10.1101/2022.03.02.480688

**Authors:** Meredith N. Frazier, Isha M. Wilson, Juno M. Krahn, Kevin John Butay, Lucas B. Dillard, Mario J. Borgnia, Robin E. Stanley

## Abstract

Coronaviruses generate double-stranded (ds) RNA intermediates during viral replication that can activate host immune sensors. To evade activation of the host pattern recognition receptor MDA5, coronaviruses employ Nsp15, which is uridine-specific endoribonuclease. Nsp15 is proposed to associate with the coronavirus replication-transcription complex within double-membrane vesicles to cleave these dsRNA intermediates. How Nsp15 recognizes and processes dsRNA is poorly understood because previous structural studies of Nsp15 have been limited to small single-stranded (ss) RNA substrates. Here we present cryo-EM structures of SARS-CoV-2 Nsp15 bound to a 52nt dsRNA. We observed that the Nsp15 hexamer forms a platform for engaging dsRNA across multiple protomers. The structures, along with site-directed mutagenesis and RNA cleavage assays revealed critical insight into dsRNA recognition and processing. To process dsRNA Nsp15 utilizes a base-flipping mechanism to properly orient the uridine within the active site for cleavage. Our findings show that Nsp15 is a distinctive endoribonuclease that can cleave both ss- and dsRNA effectively.

## Main Text

Severe acute respiratory syndrome coronavirus 2 (SARS-CoV-2) has infected millions worldwide and led to the unprecedented Covid-19 global health pandemic. Despite the rapid development of effective vaccines, new anti-viral treatments are urgently needed due to emerging SARS-CoV-2 variants and the prevalence of break-through infections amongst those already vaccinated. Coronaviruses are a family of large positive sense single stranded (ss) RNA viruses that utilize a multi-subunit replication-transcription complex (RTC) to replicate the viral genome and synthesize sub-genomic viral mRNA transcripts within double membrane vesicles ^1,2^. The RTC is composed of multiple viral non-structural proteins (Nsps), including an RNA-dependent RNA polymerase (RdRp, Nsp12), RNA helicase (Nsp13), dual exonuclease/N7-Methyltransferase (Nsp14), 2’O-Methyltransferase (Nsp16), and multiple accessory/cofactor subunits (Nsp7, Nsp8, Nsp9 and Nsp10)^2–6^. The RTC is a leading target for new anti-viral therapeutics such as nucleotide analogues that specifically target the Nsp12 polymerase subunit^6–8^.

During viral replication the RTC generates double-stranded (ds) RNA intermediates, which are well established activators of the innate immune defense system^2,9–11^. To prevent the accumulation of long dsRNAs, which activate the pattern recognition receptor MDA5, coronaviruses employ a uridine specific endoribonuclease (EndoU, Nsp15)^12,13^. EndoU like enzymes are also found across the larger nidovirus family suggesting that this endonuclease activity is critically important for single-stranded (ss) RNA viruses. However, Nsp15 is one of the most understudied non-structural proteins^5,14,15^. In virus infected cells Nsp15 has been shown to cleave the polyU tail at the 5’-end of the negative strand replication intermediate as well as numerous sites within the positive strand^12,16^. In vitro, Nsp15 has broad cleavage specificity towards ssRNA substrates and is largely guided to its cleavage sites by recognition of uridine^17^. Structures of SARS-CoV-2 Nsp15 in complex with single and di-nucleotide substrates have revealed the molecular basis for uridine specificity and provided molecular details into the RNase A-like transesterification reaction^17–19^. Recent molecular modeling suggests that beyond its role in cleaving viral RNA, Nsp15 may play a pivotal role as the central scaffold for the RTC complex^5^. Nsp15 assembles into a 240 kDa hexamer and this oligomerization is required for nuclease activity^20^. While the molecular basis for oligomerization is still not fully understood^19^, it has been proposed that the Nsp15 hexamer functions as the central scaffold for the RTC to coordinate the RTC’s numerous enzymatic activities by serving as a platform for binding dsRNA generated during replication^5^. This model is further supported by work establishing that Nsp15 co-localizes with members of the RTC^21,22^.

Earlier work with SARS-CoV-1 Nsp15 revealed that the enzyme can cleave both ss- and dsRNA substrates^14^. It is not known how Nsp15 can cleave dsRNA as this would require movement of a base-paired uridine into the uridine binding pocket, where it specifically interacts with a well conserved serine residue^17–19^. Furthermore, outside of the EndoU active site it remains unclear how the largely electronegative surface of the Nsp15 hexamer could engage a dsRNA substrate. Central to the Nsp15 RTC scaffold model is Nsp15’s ability to engage dsRNA across multiple Nsp15 promoters, however there is no experimental to support this model of dsRNA binding^5^. To address these gaps in our knowledge we solved structures of SARS-CoV-2 bound to dsRNA and determined the significance of RNA-protein interfaces by site-directed mutagenesis coupled with nuclease assays. We discovered that SARS-CoV-2 Nsp15 can process both ss- and dsRNA through a unique base-flipping mechanism.

## Results

### Structure of SARS-CoV-2 Nsp15 bound to dsRNA

To experimentally determine how SARS CoV-2 Nsp15 interacts with dsRNA, we determined a cryo-EM structure of the Nsp15 hexamer bound to a 52-nt blunt termini dsRNA^23^ (Fig. 1, table S1, Table 1 and fig. S1). To prevent RNA cleavage, we used a catalytic deficient mutant of Nsp15 (H235A) for structural studies^19^. We incubated Nsp15 with an excess of annealed dsRNA for 1 hour prior to vitrification and could visualize dsRNA extending from the Nsp15 hexamer in the 2D classes (Fig. 1A). Following 3D classification and refinement the Nsp15 dsRNA complex converged on a single asymmetric class with one dsRNA bound to the Nsp15 hexamer (fig. S1). The overall resolution of the reconstruction goes to 3.4 Å, and there is clear density for dsRNA engaged by the Nsp15 hexamer however, the local resolution is lower for the dsRNA (Fig. 1B-D). The dsRNA substrate we used contains multiple uridine nucleotides in each strand, leading to heterogeneity in the register of the dsRNA (Fig. 1B-D). Despite the nucleotide heterogeneity we were able to dock in a model of dsRNA containing 35 nucleotides in each strand and spanning a length of ~100 Å (fig. S2). The local resolution of the EndoU domains that do not engage the dsRNA is also lower supporting our earlier observation that substrate is required to fix the EndoU domain into position^19^. To improve the resolution of the dsRNA surrounding the engaged EndoU active site we collected an additional dataset to increase the number of particles, resulting in a second reconstruction with an overall resolution to 3.2 Å and improved resolution in the engaged EndoU active site (fig. S3 and Table 1). Overall, the reconstructions are very similar to one another. The first reconstruction in which we can see more overall density for the dsRNA was used for global analysis and docking of the dsRNA, while the second reconstruction was used for a specific analysis of the engaged endoU active site.

**Fig. 1.**
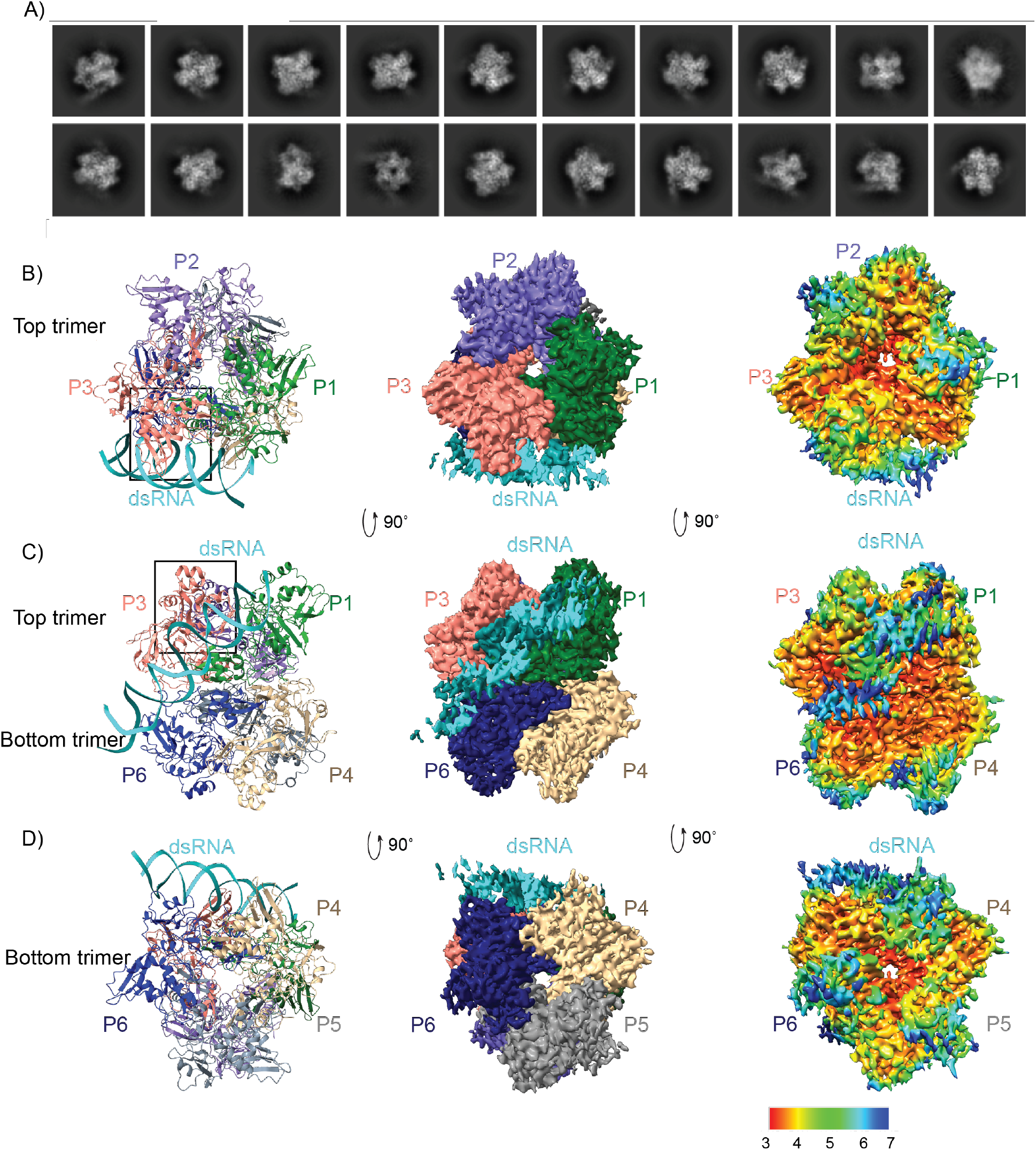
DsRNA binds Nsp15 through interactions with two platforms in addition to the EndoU active site. (**A**) Selected 2D classifications of Nsp15 bound to dsRNA. DsRNA is visible extending away from the complex. (**B**-**D**) Top, side, and bottom views of the complex in ribbon (left), EM density (middle), and local resolution (right) views.

**Table 1.** Cryo-EM collection and processing statistics.

| **Data collection and processing** | **Nsp15/dsRNA dataset 1**<br>**(EMD-26073, PDB: 7TQV)** | **Nsp15/dsRNA dataset 2**<br>**(EMD-25915 PDB: 7TJ2)** |
| --- | --- | --- |
| Microscope | Talos Arctica | Titan Krios |
| Detector | Gatan K2 Summit | Gatan K3 Bioquantum |
| Nominal magnification | 45000x | 81000x |
| Voltage (kV) | 200 | 300 |
| Electron exposure (e<sup>-</sup>/Å<sup>2</sup>) | 54 | 60 |
| Defocus range (μm) | -0.8 to -1.8 | -1.2 to 2.2 |
| Pixel size (Å) | 0.93 | 0.53 |
| Symmetry imposed | C1 | C1 |
| Number of Micrographs | 2435 | 752 |
| Initial particle images | 1,201,304 | 254,136 |
| Final particle images | 398,516 | 97,592 |
| **Refinement** | **Dataset 1** | **Combined Datasets** |
| Resolution (Å)/FSC threshold | 3.43/0.143 | 3.19/0.143 |
| B-factor used for map sharpening (Å<sup>2</sup>) | 162.7 | 171.3 |
| **Map to model CC** |  |  |
| CC (mask) | 0.69 | 0.76 |
| CC (volume) | 0.68 | 0.74 |
| CC (peaks) | 0.56 | 0.65 |
| CC (box) | 0.68 | 0.69 |
| **Model composition** |  |  |
| Non-hydrogen atoms | 17793 | 17084 |
| Protein residues | 2083 | 2007 |
| Nucleic acid | 66 | 62 |
| **Mean B factors (Å<sup>2</sup>)** |  |  |
| Protein | 85.95 | 58.77 |
| Nucleic acid | 277.22 | 150.81 |
| **R.m.s. deviations** |  |  |
| Bond lengths (Å) | 0.002 | 0.002 |
| Bond angles (°) | 0.383 | 0.370 |
| **Validation** |  |  |
| Molprobity score | 1.52 | 1.39 |
| Clashscore | 5.00 | 3.74 |
| Poor rotamers (%) | 0.82 | 1.64 |
| **Ramachandran plot** |  |  |
| Favored (%) | 96.18 | 97.69 |
| Allowed (%) | 3.82 | 2.31 |
| Disallowed (%) | 0.00 | 0.00 |

### Nsp15 engages dsRNA through multiple interfaces

In both cryo-EM reconstructions the Nsp15 hexamer asymmetrically engages one dsRNA substrate. Nsp15 forms a hexamer of back-to-back trimers. The dsRNA interacts with three of the six Nsp15 protomers; two from the top trimer (P1 and P3), and one from the bottom trimer (P6, Fig. 1B-D). Furthermore, across the three interacting protomers the dsRNA ultimately interacts with residues spanning all three domains that comprise an Nsp15 protomer (N-terminal, Middle, and catalytic EndoU domain; Fig. 2, fig. S4). We identified 4 key RNA-protein interfaces within the structure (Fig 2A, fig. S4). The N-terminal domain (NTD) of Nsp15, including residues Q19 and Q20, interacts with the dsRNA through P1 in the top trimer (P1-B). Two different interfaces stem from the middle domain (MD): P1 in the top trimer (P1-A), and P6 in the bottom trimer, both of which predominantly interact with the backbone through the major groove. Key residues in these interfaces include K65, K111, and K150. Finally, the EndoU domain of P3 in the top trimer interacts with the dsRNA through interactions with both the minor and major groove. The minor groove interaction is mediated by EndoU residues such as, H243 and Q245 while the major groove interaction includes residues W333, Y343, and E340. The structure revealed that Nsp15 interacts with both strands of the dsRNA, and except for the active site uridine, all of the protein-RNA interactions are RNA backbone mediated. This supports earlier work demonstrating that Nsp15 has broad specificity towards uridine containing substrates^16,17^.

**Fig. 2.**
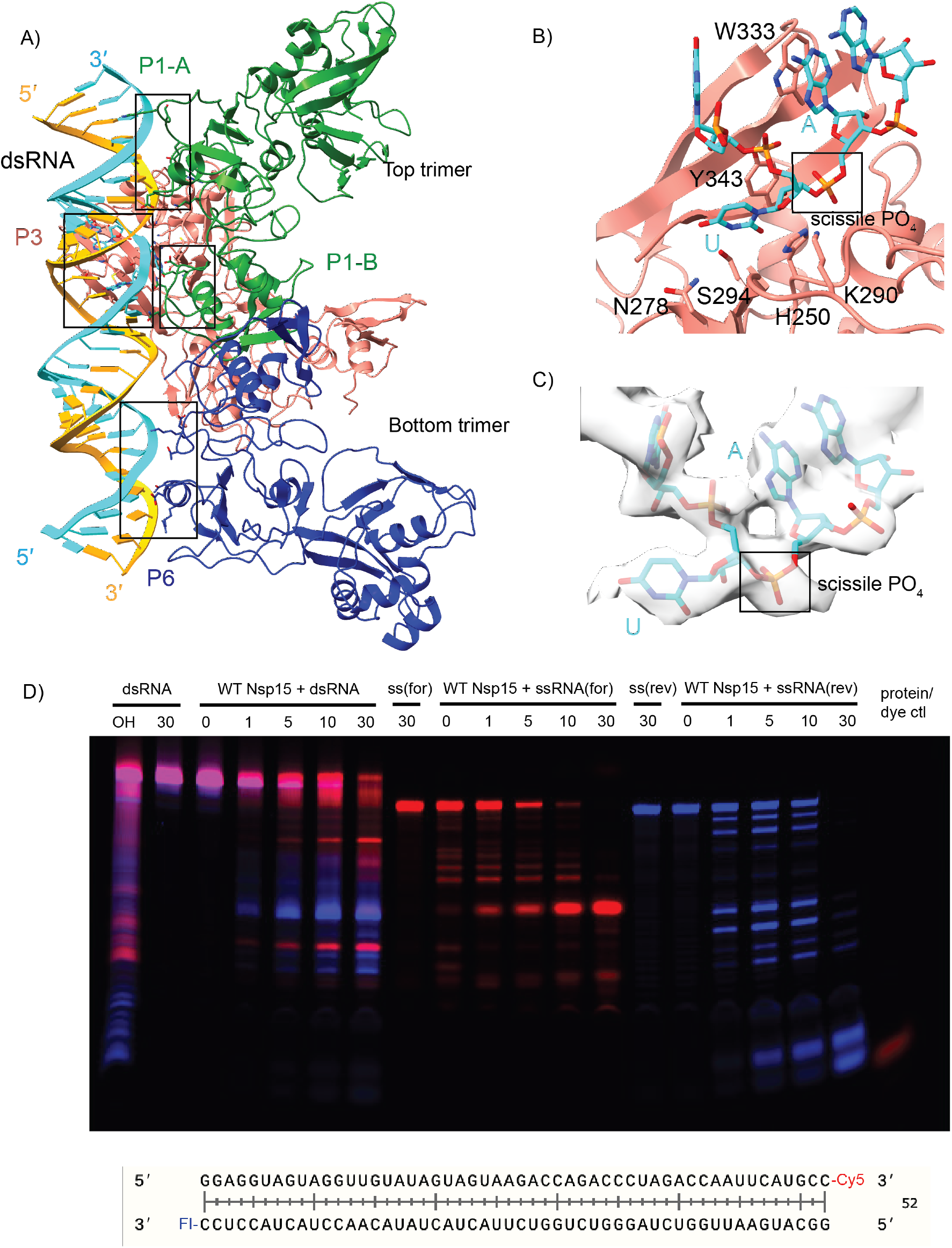
Nsp15 can cleave both ss- and dsRNA. (**A**) DsRNA interacts with three Nsp15 protomers, across both trimers. P1 and P6 form “platforms” that support RNA cleavage by P3. (**B**) Close-up of the active site (P3). Critical residues are shown in stick format. Uridine flips out to interact with S294 and N278, which provides optimal positioning for the catalytic triad. The 3’ base is stabilized by W333. (**C**) Cryo-EM density for the RNA engaged in the active site. (**D**) Time-course cleavage reaction for the dsRNA as well as each strand alone. Nsp15 cleaves ssRNA more quickly than dsRNA, and prefers different positions depending on that context. OH: alkaline hydrolysis of dsRNA. ssRNA(for) is the Cy5 labeled strand (red); ssRNA(rev) is the FI-labeled strand (blue).

Overall, our structures are in good agreement with the predicted position of dsRNA binding interfaces derived from the computational model of Nsp15 proposed in the RTC Nsp15 based scaffold(fig. S5)^5^. However, unlike the model which proposes multiple copies of RNA bound to the Nsp15 hexamer, the majority of our cryo-EM 2D classes revealed only one RNA bound per hexamer (Fig 1). Previously determined cryo-EM structures of Nsp15 determined in the presence and absence of small RNA substrates have revealed both symmetric and asymmetric states, thus we assume it is possible for Nsp15 to engage more than one dsRNA substrate at a time^17,19^. We hypothesize that the Nsp15 scaffold is a very transient part of the RTC as this platform positions the dsRNA within the EndoU active site. This Nsp15-scaffolded RTC may only form during certain phases of replication (i.e. discontinuous transcription). Alternatively, the assembly of the RTC complex with Nsp15 in the center could lock Nsp15 into an inactive conformation incapable of cleaving dsRNA.

### Nsp15 base flipping mechanism

While the register for the RNA is ambiguous in our structure, we observed density for a nucleotide engaged in the active site, which we modeled as a uridine. The engaged uridine flips out from the dsRNA helix and is positioned in the active site as previously observed in UMP, and di-nucleotide bound structures (fig. S6)^17–19^. As a result of the flipping of the engaged uridine the base 3’ of the uridine, which we modeled as an A, is destabilized from the helix. In our structure this base engages with W333, which likely stabilizes that base to support cleavage of the phosphate backbone 3’ of the engaged uridine (Fig. 2B). Collectively both the flipping of the uridine and the engagement the 3’ base with W333 re-position the scissile phosphate within the Nsp15 active site composed of the catalytic residues H235, H250, and K290. Aside from the flipped base, we do not have sufficient density to determine if the global dsRNA structure is otherwise perturbed. Many nucleic acid processing enzymes utilize a base-flipping mechanism to position substrates within their active sites, however most of these enzymes do not flip bases from normal, A-form dsRNA^24–29^. The base-flipping mechanism in Nsp15 is reminiscent to the mechanism used by adenosine deaminases acting on RNA (ADARs), which convert adenosine to inosine in dsRNA. Like Nsp15, ADARs, use a base flipping mechanism to position the reactive adenosine within the editing active site. Thus, Nsp15 appears to join a small group of RNA enzymes, including ADARS, known to flip a base from a normal duplex RNA substrate^24^. It remains to be seen if Nsp15 actively flips out the engaged uridine or if it stabilizes the flipped-out base through a passive process.

### Nsp15 cleaves ss- and dsRNA

Following structure determination, we confirmed that SARS-CoV-2 Nsp15 cleaves dsRNA substrates. Previous studies with SARS-CoV-1 and human coronavirus 299E (HCov-229E) Nsp15 using a 22-nt RNA substrate containing a GUU sequence suggested that Nsp15 cleaves dsRNA more efficiently than ssRNA^14^. To confirm that SARS-CoV2 Nsp15 can cleave dsRNA we carried out a time course using the 52-nt dsRNA from the cryo-EM structure with labels on the 3’ ends. We observed that Nsp15 can cleave the duplex 52-nt substrate as well as the individual strands at numerous positions. (Fig. 2D). We observed that the accumulation of cleavage products is different suggesting that the cleavage specificity for ssRNA vs dsRNA varies. A similar observation of altered specificity was previously made with SARS-CoV-1 and HCov-299E Nsp15^14^. We hypothesize that this could be a result of the base flipping mechanism. Uridines followed by strong base pairs may not be as accessible to the EndoU active site. This is supported by RNA sequencing data from mouse hepatitis virus (MHV) infected cells which showed a strong preference for adenines 3’ of Nsp15 cleavage sites detected in the positive strand^16^. Similarly, ADARs have a nearest neighbor preference for specific base-pairs before and after the reactive adenine that flips into the active site^24,30^.

To further probe Nsp15’s ability to cleave dsRNA we measured cleavage of an alternative dsRNA substrate. Our structure suggests that dsRNA of approximately 35-nts is sufficient to interact across the Nsp15 hexamer, so we expanded on the nucleocapsid protein transcriptional regulatory sequence (TRS-N) previously used to characterize ssRNA cleavage preferences (table S1, ^17^). TRS sequences are found upstream in many of the viral subgenomic RNAs, are important for facilitating discontinuous transcription and are known to form dsRNA intermediates ^2,31^. To compare nuclease activity, a 60 min time course reaction with 35-nt ss- and dsRNA was performed and the disappearance of the uncleaved RNA band quantified (Fig. 3-4, fig. S7-S8). Similar to our results with the 52-nt dsRNA substrate, Nsp15 can cleave both the ss- and dsRNA TRS and we observe that the ssRNA is cleaved faster than the dsRNA under the same reaction conditions. We hypothesize that ssRNA is cleaved faster because it does not have to undergo base-flipping to be accessible to the Nsp15 EndoU active site. We also observe different cleavage patterns for the ss and ds TRS (fig. S7-8) further suggesting that Nsp15’s substrate specificity is altered for ss and dsRNA.

**Fig. 3.**
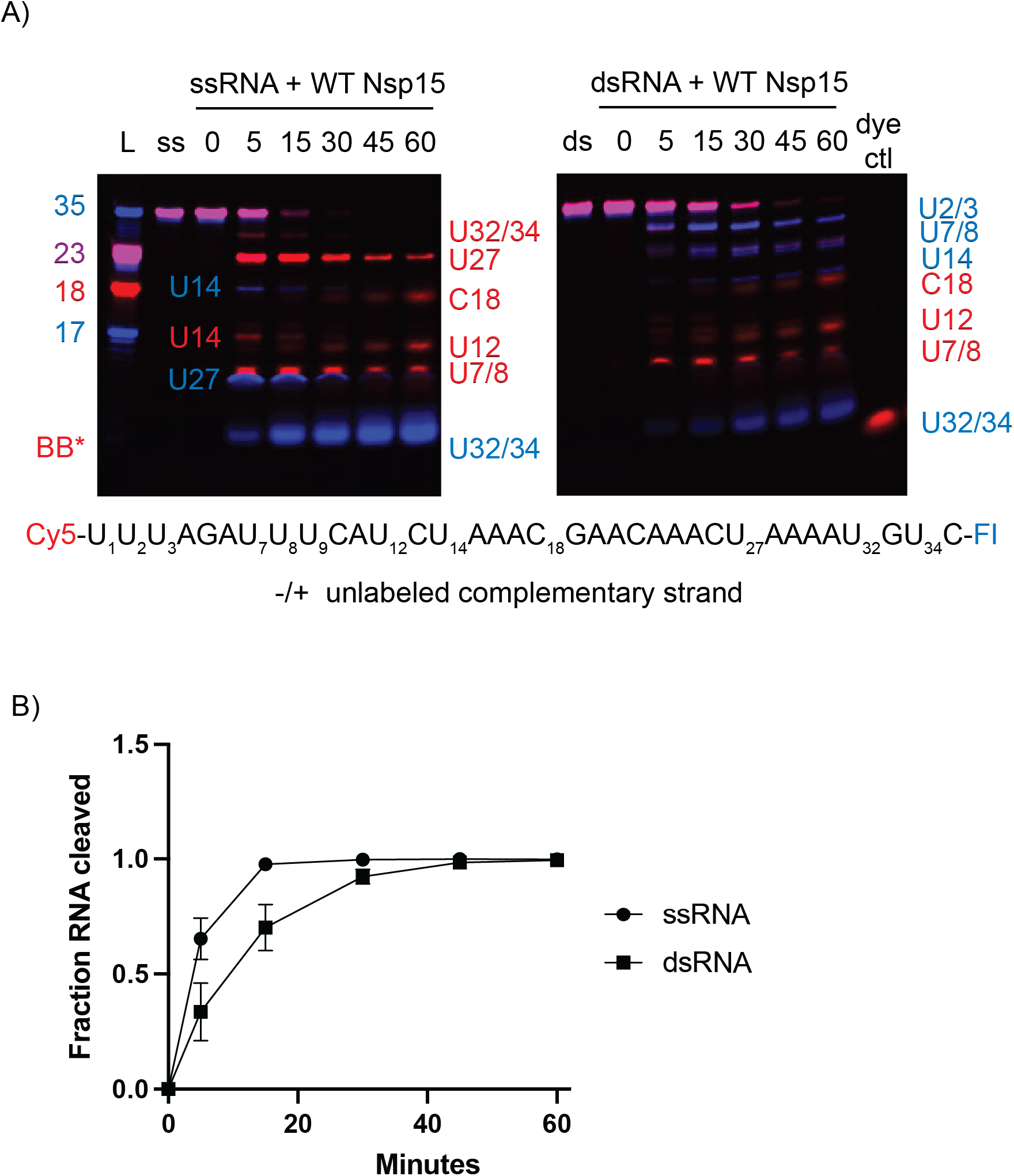
Comparison of WT Nsp15 activity on ss- and ds-RNA. (**A**) Time course reactions with WT Nsp15 were performed on double-labeled ssRNA +/- the unlabeled complementary strand. Cleavage products are labeled to the right of the gel, with the color corresponding to which product (Cy5 or FI) it is. (**B**) RNA cleaved was quantified by disappearance of the uncleaved RNA band, normalized to the 0 time point for that reaction. The average and standard deviation for at least three independent reactions are graphed.

**Fig. 4.**
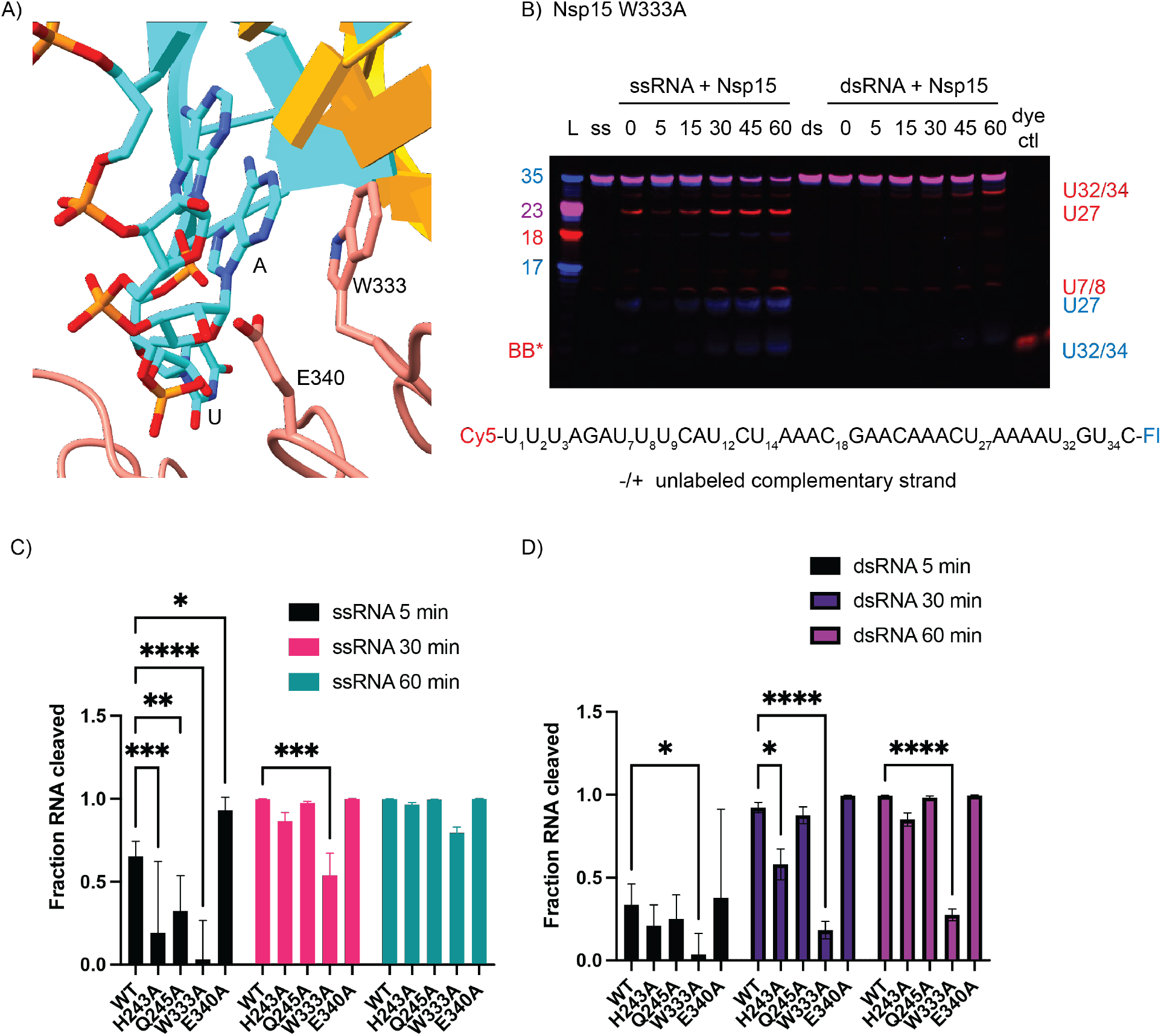
EndoU domain mutations in Nsp15 affect dsRNA cleavage. (**A**) Zoomed in depiction of the active site mutants, focused on W333. (**B**) Time course reactions for Nsp15 active site mutants were performed with double-labeled ssRNA +/- the unlabeled complementary strand. A representative time course is shown for Nsp15 W333A. Cleavage products are labeled to the right of the gel, with the color corresponding to which product (Cy5 or FI) it is. (**C-D**) RNA cleaved was quantified by disappearance of the uncleaved RNA band, normalized to the 0 time point for that reaction. The average and standard deviation for at least three independent reactions are graphed.

### W333 mediates dsRNA cleavage

To test the importance of dsRNA-interacting residues identified by our structure, we made point mutants for several residues spanning the three domains of Nsp15 (table S2, and fig. S9). We compared the activity of these mutants on both 35-nt ss- and dsRNA using gel-based cleavage assays (Fig. 4, fig. S7-8). Most of the single point mutants exhibited little to no defects in ssRNA and small defects in dsRNA processing, likely because of the large number of residues that support dsRNA engagement. One mutant, E340A gave rise to an increase in cleavage activity (fig. S8). Given that RNA is negatively charged, the positioning of E340 in the active site may be important for regulating nuclease activity. We also identified a critical active site mutation that is required for dsRNA cleavage. Our structure suggests that W333 plays a crucial role in stabilizing the dsRNA, with a flipped-out base by interacting with the base 3’ of the cleavage site. While W333A results in a 20% reduction of ssRNA cleavage at 60 min, the defect in dsRNA is much more severe, resulting in 75% less cleavage at 60 min (Fig. 4). This observation is further supported by our earlier structure of Nsp15 bound to ssRNA in the pre-cleavage state, in which we could not observe strong density for the base 3’ of the cleavage site ^17^. Collectively this suggests that W333 is a critical regulator for stabilizing the base 3’ of the cleavage site in dsRNA but is not essential for positioning the same base within a flexible ssRNA substrate (Fig. 5).

**Fig. 5.**
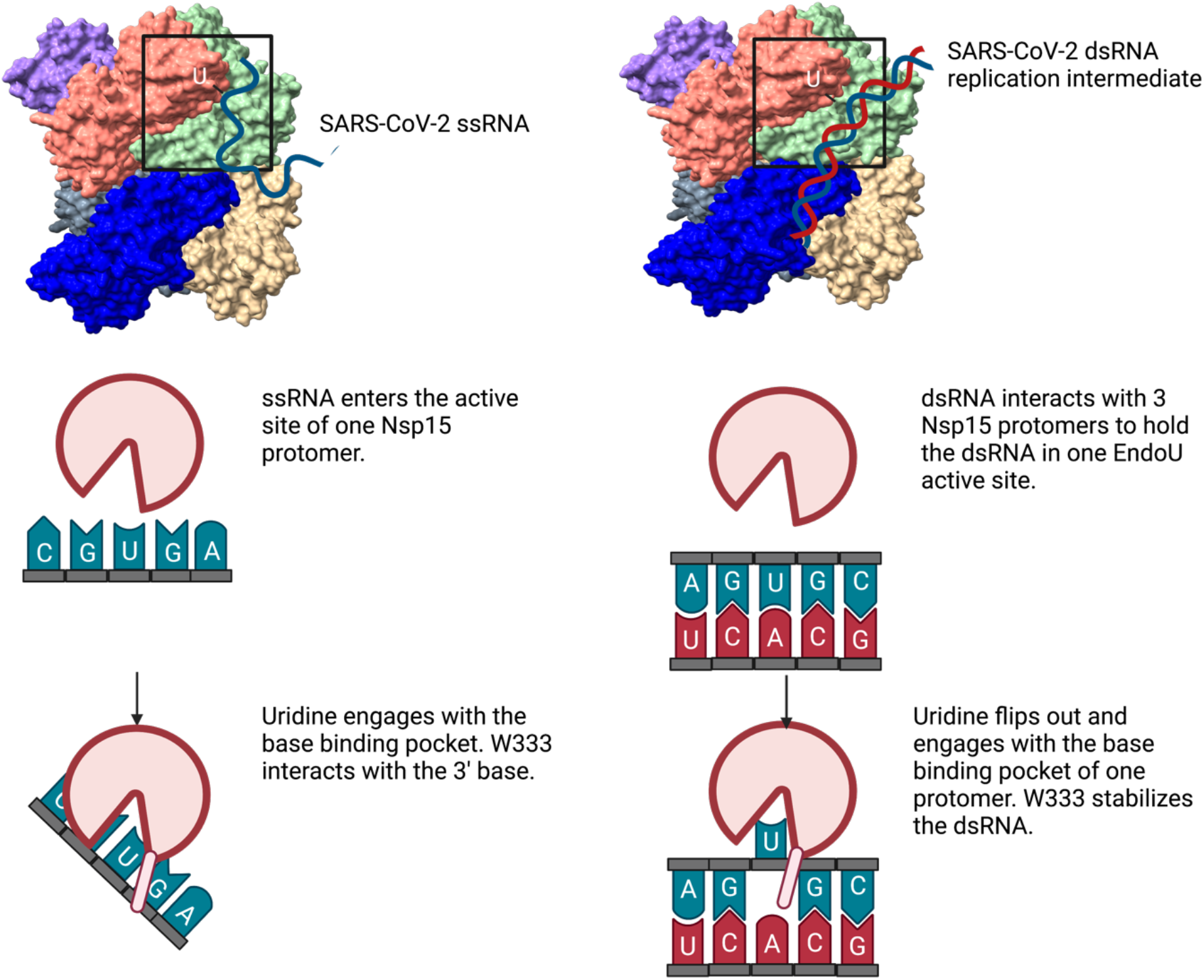
Model comparing ss- and dsRNA cleavage by Nsp15. Nsp15 cleaves ssRNA mainly through engagement of the U in the active site, with limited binding by upstream or downstream nucleotides. ssRNA is readily accessible to the Nsp15 active site. Nsp15 cleaves dsRNA by multiple interaction sites, spanning three protomers and both trimers that compose the hexamer. The U base must flip out of the base-paired helix to be cleaved; distortion in the RNA is stabilized through pi-stacking with W333. Figure made in Biorender.

## Discussion

Overall, our structures and biochemical assays provide molecular insight into the mechanism of dsRNA processing by SARS-CoV-2 Nsp15. We observed that the Nsp15 hexamer forms a platform across multiple protomers to engage dsRNA. Three of the six protomers of Nsp15 form an electropositive surface for dsRNA of approximately 100 Å in length. There are numerous ways that ribonucleases and other RNA binding proteins interact with RNA but many of these interactions are mediated by small RNA binding domains^32^. For example, the small structurally conserved dsRNA Binding Domain (dsRBD) is utilized by many RNA processing enzymes either on its own or in combination with other domains to mediate dsRNA binding^33^. This domain has been well-studied in RNase III family members, such as dicer and drosha, where it has been shown to specifically recognize features within the minor groove of A-form RNA^34^.

ADARs also contain multiple dsRBDs, outside of the catalytic deaminase domain, that contribute to RNA binding and are required for editing of some dsRNA substrates^35^. In contrast Nsp15 does not use a single domain or even a single protomer to bind RNA, rather it uses all three of its domains across multiple protomers to engage dsRNA. This unique RNA interface provides the molecular basis for Nsp15’s dependence on oligomerization. Our study reveals that the oligomerization of Nsp15 is necessary to form the platform for RNA binding with contributions from three of the six protomers.

Studies with Nsp15 from other coronaviruses, as well as Nsp11, the EndoU containing Nsp in arteriviruses, have demonstrated oligomerization is necessary for endoribonuclease activity although the type of oligomer (dimer, hexamer) varies across viruses^36,37^. We compared our Nsp15/dsRNA structure with previous structural work that assessed the role of oligomerization and activity of nidovirus EndoU enzymes (fig. S10). Structures of MERS-CoV and SARS-CoV-1 Nsp15 superpose very well with SARS-CoV-2 Nsp15^37^, suggesting that the dsRNA platform is conserved across CoVs. In contrast to Nsp15, Porcine Reproductive and Respiratory Syndrome Virus (PRRSV) Nsp11 forms a dimer, not a hexamer. We modeled dsRNA onto the PRRSV Nsp11 dimer by superposition of the EndoU domains and observed that the dsRNA fits in the groove formed between Nsp11 protomers. The π-stacking tryptophan is conserved and positioned to stabilize the dsRNA when the uridine is flipped out. This observation suggests that the base-flipping mechanism is conserved across nidovirus EndoU enzymes while the dsRNA platform differs.

Beyond facilitating dsRNA binding and cleavage, the Nsp15 RNA platform may further support the viral RTC complex to coordinate replication and transcription. Molecular modeling suggests that Nsp15 could be the central scaffold for the entire RTC complex^5^. Nsp15 has been proposed to be the central platform that coordinates the formation of an RTC superstructure including 6 copies of Nsp11, Nsp12, Nsp13, Nsp14, Nsp15, and Nsp16. This superstructure provides a putative mechanism for the movement of the newly transcribed RNA into the capping and nuclease active sites within the RTC^5^. While experimental work is needed to confirm the formation of the RTC superstructure our structures reveal that the extensive dsRNA binding interface within the Nsp15 hexamer could support the RTC. However, we hypothesize that the RTC superstructure forms very transiently otherwise the Nsp15 EndoU active sites would likely degrade a significant portion of the viral RNA.

Our work further establishes how Nsp15 cleaves dsRNA, broadening the repertoire of Nsp15 cleavage activity against a variety of RNA substrates (Fig. 5). From earlier RNA bound structures of Nsp15 it was unclear how dsRNA would be able to access the EndoU active site. Flexible ssRNA substrates are positioned into the active site through uridine recognition in the uridine binding pocket. Our structure and mutagenesis revealed that Nsp15 can support dsRNA cleavage through a base flipping mechanism that positions the engaged uridine within the uridine binding pocket (Fig. 5). Nsp15 also utilizes a conserved tryptophan residue to position the base 3’ of the uridine. The combined action of flipping the uridine and stabilizing the 3’ base moves the phosphodiester bond following the engaged uridine into the Nsp15 active site, where is undergoes an RNase A-like transesterification reaction. Through this unique mechanism Nsp15 can process both ss- and dsRNA substrates. The ability to cleave structurally diverse RNA substrates under the same conditions is rare in endoribonucleases, and points towards the evolution of Nsp15’s activity to be able to process both ss- and ds-viral RNA to regulate the overall level in infected cells. Bovine viral diarrhea virus, a pestivirus, also encodes for an endoribonuclease that can process ss- and ds-viral RNAs, further emphasizing the importance of viral enzymes to be multi-functional given their small genomes^38^.

Finally, Nsp15 is a promising anti-viral target, and this work reveals multiple RNA interfaces that could be targeted for structural based drug design. Nsp15 antagonizes the host immune system by limiting the accumulation of dsRNA intermediates that form during viral replication^12,13,15,39,40^. Inactivating Nsp15 in porcine epidemic diarrhea virus has been shown to elicit a strong immune response in infected animals and reduced viral shedding and mortality^41,42^. Our work establishes that Nsp15 contains four previously unknown RNA binding interfaces that support dsRNA engagement, and further emphasizes that blocking Nsp15 oligomerization could have therapeutic benefit.

## Acknowledgments

We would like to thank the NIEHS Mass Spectrometry Research and Support Group, especially Jason Williams, for protein identification analyses. We would like to thank Drs. Traci Hall and Marcos Morgan for their critical review of this manuscript. We would like to thank all members of the NIEHS Molecular Microscopy Consortium for their help with data collection. We would like to thank Dr. Rick Huang and Allison Zeher for help with cryo-EM data collection. This work utilized the NCI/NICE Cryo-EM Facility. This work was supported by the US National Institutes of Health Intramural Research Program; US National Institute of Environmental Health Sciences (NIEHS, NIEHS/NIH ZIA ES103247(RES); NIEHS/NIH ZIC ES103326 (MJB)). This work was also supported by the NIH Intramural Targeted Anti-COVID-19 (ITAC) Program funded by the National Institute of Allergy and Infectious Diseases (NIAID, NIAID/NIH 1ZIAES103340 (RES)).

## Author contributions

Conceptualization: MNF, RES; Methodology: MNF, JMK, IMW, KJB, LBD, MJB, RES; Investigation: MNF, JMK, IMW, KJB, LBD, MJB, RES; Visualization: MNF, JMK, IMW, KJB, LBD, MJB, RES; Funding acquisition: MJB, RES; Writing – original draft: MNF, RES; Writing – review & editing: MNF, JMK, IMW, KJB, LBD, MJB, RES

## Data and materials availability

The cryo-EM structures have been deposited in the PDB and EMDB with the following accession codes: EMD-25915, PDB: 7TJ2, EMD-26073, PDB: 7TQV. All other data presented in this manuscript are available in the main text or the supplementary materials.

