## Supplementary Information for "Flipped Over U: Structural Basis for dsRNA Cleavage by the SARS-CoV-2 Endoribonuclease"

**Affiliations:**
[1]Signal Transduction Laboratory, National Institute of Environmental Health Sciences, National Institutes of Health, Department of Health and Human Services, 111 T. W. Alexander Drive, Research Triangle Park, NC 27709, USA

[2]Genome Integrity and Structural Biology Laboratory, National Institute of Environmental Health Sciences, National Institutes of Health, Department of Health and Human Services, 111 T. W. Alexander Drive, Research Triangle Park, NC 27709, USA

[3]Present Address: Program in Molecular Biophysics, Johns Hopkins University, 3400 N Charles St. Baltimore, MD 21218

**This File Includes:**

Materials and Methods

Supplemental Figures 1-10

Supplementary Tables 1-2

### Methods

#### Protein expression and purification

Wild type (WT) and mutant Nsp15 constructs were created as described previously[1,2]. Nsp15 was overexpressed in *E. coli* C41 (DE3) competent cells in Terrific Broth with 100 mg/L ampicillin. At an optical density (600 nm) between 0.8-1.0, cultures were cooled prior to induction with 0.2 mM Isopropyl β-D-1-thiogalactopyranoside (IPTG). Cells were harvested after overnight expression at 16˚C and stored at -80˚C until use. Nsp15 purification was done as described previously[1,2]. Briefly, cells were resuspended in Lysis Buffer (50 mM Tris pH 8.0, 500 mM NaCl, 5% glycerol, 5 mM β-ME, 5 mM imidazole) supplemented with cOmplete EDTA-free protease inhibitor tablets (Roche) and disrupted by sonication. The lysate was clarified at 26,915 x g for 50 minutes at 4˚C and then incubated with TALON metal affinity resin (Clontech). His-Nsp15 was eluted from the resin with 250 mM imidazole, and buffer exchanged into Thrombin Cleavage Buffer (50 mM Tris pH 8.0, 150 mM NaCl, 5% glycerol, 2 mM β-ME, 2 mM $CaCl_2$) for cleavage at room temperature for 4 hours. The cleavage reaction was repassed over TALON resin and quenched with 1 mM phenylmethylsulfonyl fluoride (PMSF) prior to gel filtration using a Superdex-200 column equilibrated in SEC buffer (20 mM Hepes pH 7.5, 150 mM NaCl, 5 mM $MnCl_2$, 5 mM β-ME).

#### Cryo-EM sample preparation

RNA oligos were annealed at 250 uM final concentration by incubating equimolar amounts for 5 min at 75˚C and then cooled on the heat block for 1.5 hours. Purified Nsp15 H235A was diluted in a low-salt buffer (20 mM Hepes pH 7.5, 100 mM NaCl, 5 mM $MnCl_2$, 5 mM β-ME) to 0.75 µM and incubated with excess RNA substrate (50 µM) for 1 hour at 4˚C. The purified protein was plunge frozen on homemade customized support grids made of C-flat R1.2/1.3 (Protochips) sputtered with 30nm thick layer of gold on the grid bar side using Leica ACE-600 sputterer. Before specimen deposition, the grids were pretreated on a plasma cleaner (Tergeo, model) in immersion mode with a power of 38W for a period of 75s. The Nsp15/RNA mixture (3 µL) was deposited onto the grids inside the chamber of a Leica EM-GP2 vitrification robot held at 15˚C and RH of 90% humidity. The grid was back-blotted for 3 seconds (Whatman Grade 40 filter paper), and then quickly plunged into liquid ethane kept at 90K. The vitrified sample on the grid was then transferred to liquid nitrogen for storage.

#### Data collection and processing

Nsp15 images were collected using a Talos Arctica electron microscope at 200 keV with a Gatan K2 Summit detector, and a Titan Krios at 300 keV with a K3 Bioquantum detector. Beam-induced motion and drift were corrected using MotionCor2 [3] through Scipion 3 [4]. CryoSPARC v2 [5] was used in all subsequent image processing. The aligned dose-weighted images were used to calculate CTF parameters using CTFFIND4 [6]. For the first dataset, particles were selected using blob and template-based particle picking, downsampled by a factor of 4, extracted with a box size of 64 and subjected to an initial round of 2D classification. Full resolution particle projections from good classes were re-extracted using a box size of 256. *Ab initio* reconstruction was used to an initial model, followed by several rounds of 3D classification with localized CTF refinement and per particle motion correction. No symmetry was applied.

For the combined Arctica/Krios datasets, the Topaz Extract machine learning algorithm was used to pick particles. The extracted particles were curated and segregated using a combination of 2D classifications, Ab-Initio Model Reconstruction and Heterogeneous Refinement. The curated particles stacks from both Talos Arctica and Titan Krios were then combined. The combined particle stack that yielded the most ideal 3D map was then further refined using unsymmetrized homogeneous, Global CTF, and Non-Uniform Refinements. Maps were re-scaled to optimize RMS fit to core domain residues of reference structure PDBID 6WLC[7].

**Model building**

A SARS-CoV-2 Nsp15 cryo-EM structure (PDBID 7K0R) and model dsRNA (PDBID 6BU9) were used as starting models and fit into the cryo-EM maps using rigid body docking in Phenix[8]. A combination of rigid body and real-space refinement in Phenix as well as iterative rounds of building in COOT[9] were used to improve the fit of the models. Given the weaker density of the RNA outside the active site, it was restrained to model RNA parameters during refinement. Molprobity[10] was used to evaluate the model (Table 1). Figures were prepared using Chimera[11] and Chimera X[12].

**Urea-PAGE endoribonuclease assay**

DsRNA was prepared by incubating equimolar concentrations of complementary oligos at 75˚C for 5 minutes, followed by cooling on the heat block for 1.5 hrs. To remove any remaining unpaired RNA, samples were purified on a 20% native polyacrylamide gel. The dsRNA band was excised, and the RNA was eluted overnight in 10 mM Hepes pH 7.5, 50 mM NaCl, 0.5 u/µL RNAsin. Double fluorescently-labeled RNA substrates (5′-FI and 3′-Cy5, 500 nM) were incubated with Nsp15 (50 nM) in RNA cleavage buffer (20 mM Hepes pH 7.5, 150 mM NaCl, 5 mM $MnCl_2$, 5 mM DTT, 1 u/µL RNasin ribonuclease inhibitor) at room temperature for 30 minutes, with samples collected at 0, 1, 5, 10, and 30 minutes. The reaction was quenched with 2x urea loading buffer (8M urea, 20 mM Tris pH 8.0, 1 mM EDTA). At least three independent reactions were performed with protein from at least 2 different purifications. Due to the expected size of cleavage products and the size of bromophenol blue, loading buffer without dye was used. To monitor the gel front, a control lane of protein only with bromophenol blue was run. A ladder composed of double and single labeled oligos of different sizes were used to help assign cleavage product identities. Cy5-labeled products run anomalously compared to FI-labeled products[13,14]. The cleavage reactions were separated using 15% TBE-urea PAGE gels and visualized with a Typhoon RGB imager (Amersham) using Cy2 ($\lambda_{ex}$=488 nm, $\lambda_{em}$=515-535 nm) and Cy5 ($\lambda_{ex}$=635 nm, $\lambda_{em}$=655-685 nm) channels. RNA cleavage was measured by disappearance of the uncleaved RNA, and quantified using Image Studio Lite (LI-COR). Prism (Graphpad) was used to calculate significant differences using Dunnett's T3 multiple corrections test.

**Supplementary Figures and Tables**

**Supplementary Table 1. List of RNA oligos used in this study. All oligos were synthesized by Dharmacon (Horizon).**

| **Oligo Name** | **Oligo Sequence** | **Purpose** | **Length (nt)** |
|---|---|---|---|
| dsRNA_BB_f | GGAGGUAGUAGGUUGUAUAGUAGUAA GACCAGACCCUAGACCAAUUCAUGCC-Cy5 | Cryo-EM, cleavage assays | 52 |
| dsRNA_BB_r | GGCAUGAAUUGGUCUAGGGUCUGGUC UUACUACUAUACAACCUACUACCUCC-Fl | Cryo-EM, cleavage assays | 52 |
| DS10 | Cy5-UUUAGAUUUCAUCUAAACGAACAAAC UAAAAUGUC-FI | Cleavage assays | 35 |
| DS11 | GACAUUUUAGUUUGUUCGUUUAGAUG AAAUCUAAA | Cleavage assays | 35 |

**Supplementary Table 2. Nsp15 constructs used in this study. All mutations were synthesized by Genscript in the WT-Nsp15 backbone.**

| **Construct** | **Mutation location (domain level)** | **First created** |
|---|---|---|
| WT-Nsp15 (6xHis-thrombin-TEV/pet14-b) | N/A | Pillon et al[1] |
| Nsp15 Q19A | NTD | This study |
| Nsp15 Q20A | NTD | This study |
| Nsp15 K65A | NTD | This study |
| Nsp15 H235A | EndoU | Pillon et al[1] |
| Nsp15 H243A | EndoU | This study |
| Nsp15 Q245A | EndoU | This study |
| Nsp15 W333A | EndoU | Frazier et al[2] |
| Nsp15 E340A | EndoU | This study |

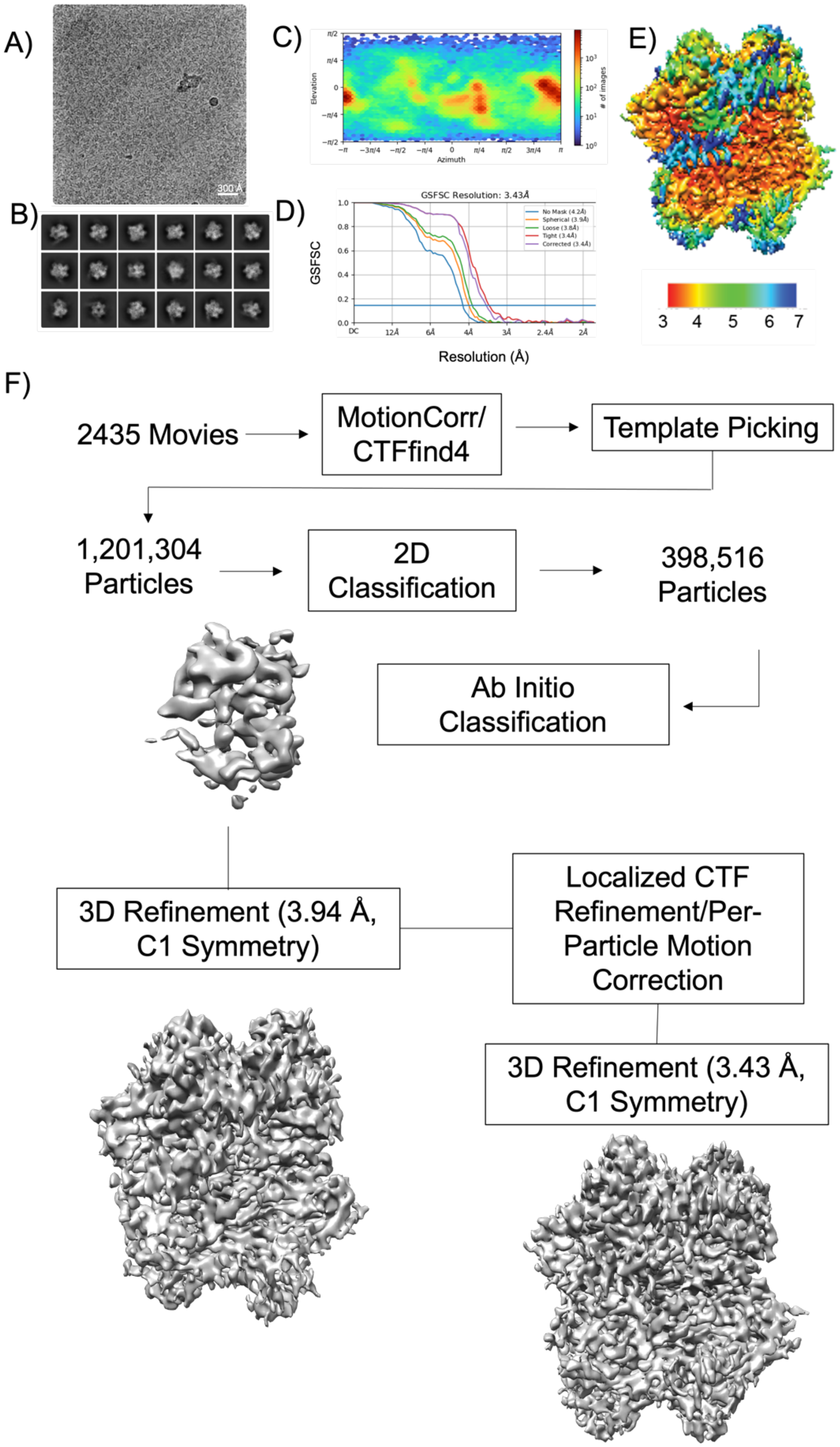


**Supplementary Figure 1.** Cryo-EM processing workflow. (A) A representative micrograph of Nsp15 H235A with excess 52-nt dsRNA in vitreous ice and (B) selected 2D classes generated from 738 movies collected from an UltrAuFoil R1.2/1.3 300 mesh grid. (C) Angular distribution of particles. (D) Fourier shell correlation (FSC) curve. The overall resolution is 3.43 Å according to the FSC 0.143 criteria[15]. (E) Cryo-EM reconstruction colored based on local resolution calculated using cryoSparc v2 [5]. (F) Cryo-EM processing workflow: picked particles (944,302) were subjected to 2D classification, 3D classification, and refinement in cryoSPARC v2 [5].

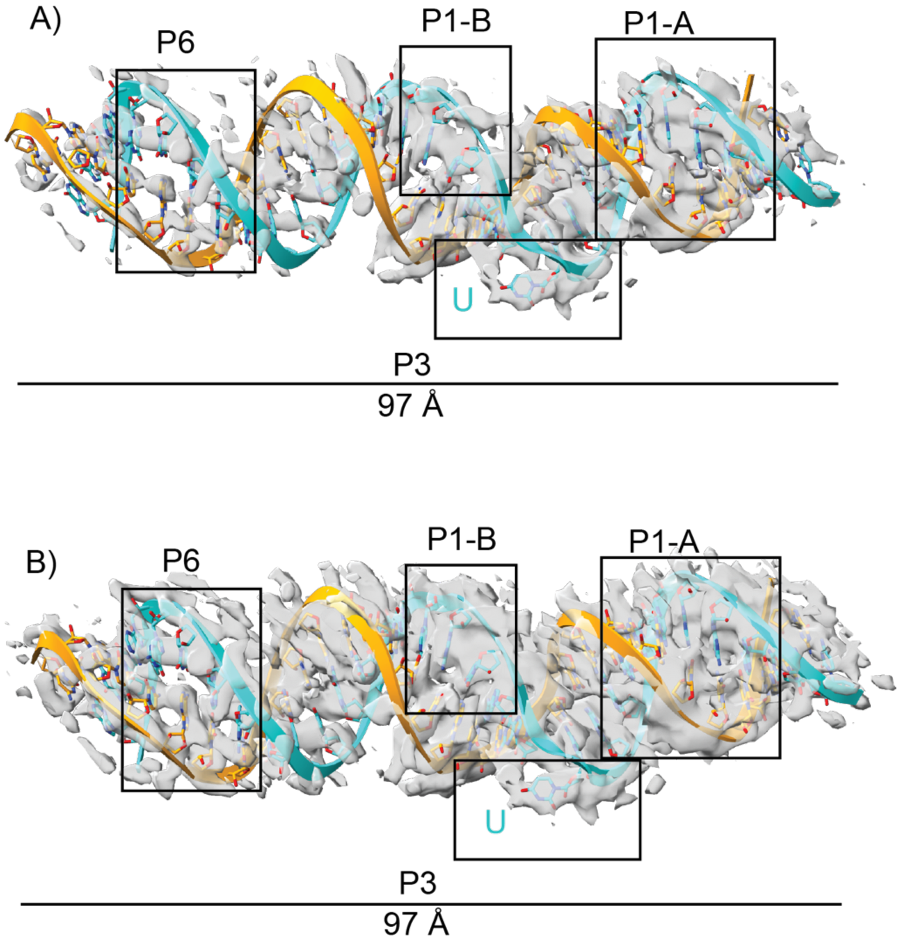


**Supplementary Figure 2.** Dataset 1 dsRNA density at two different contour levels: **A)** 0.45, **B)**, 0.36. Interaction sites are labeled with black boxes.

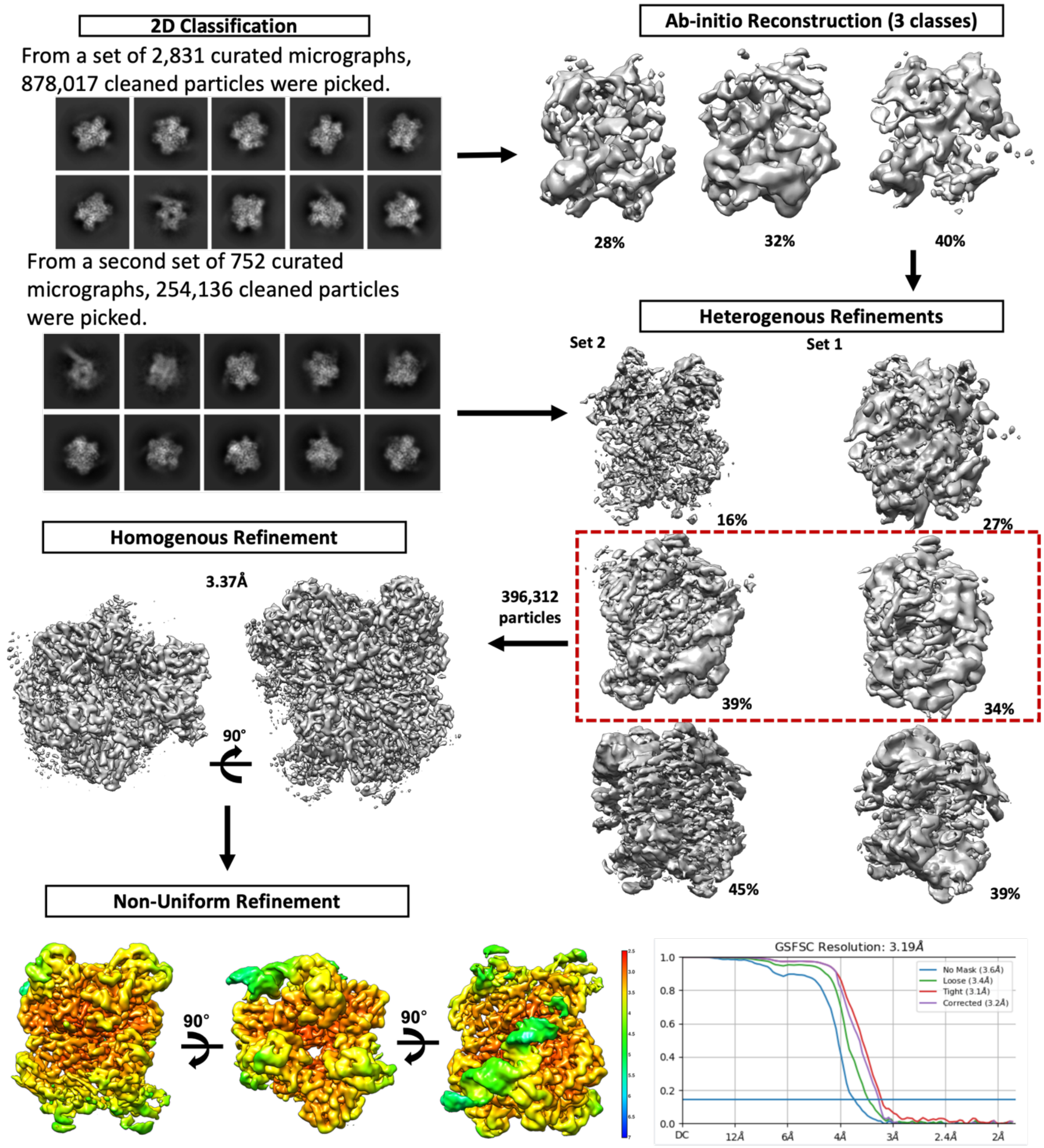


**Supplementary Figure 3.** Cryo-EM workflow for the combined Krios and Arctica datasets starting from curated 2D classes. The curated Arctica classes were used for *ab initio* reconstruction, which then underwent heterogeneous refinement with the Krios dataset curated particles inputted. The best class from the Krios refinement and the Arctica refinement were then combined for homogeneous refinement, followed by non-uniform refinement, resulting in a map at 3.19 Å global resolution.

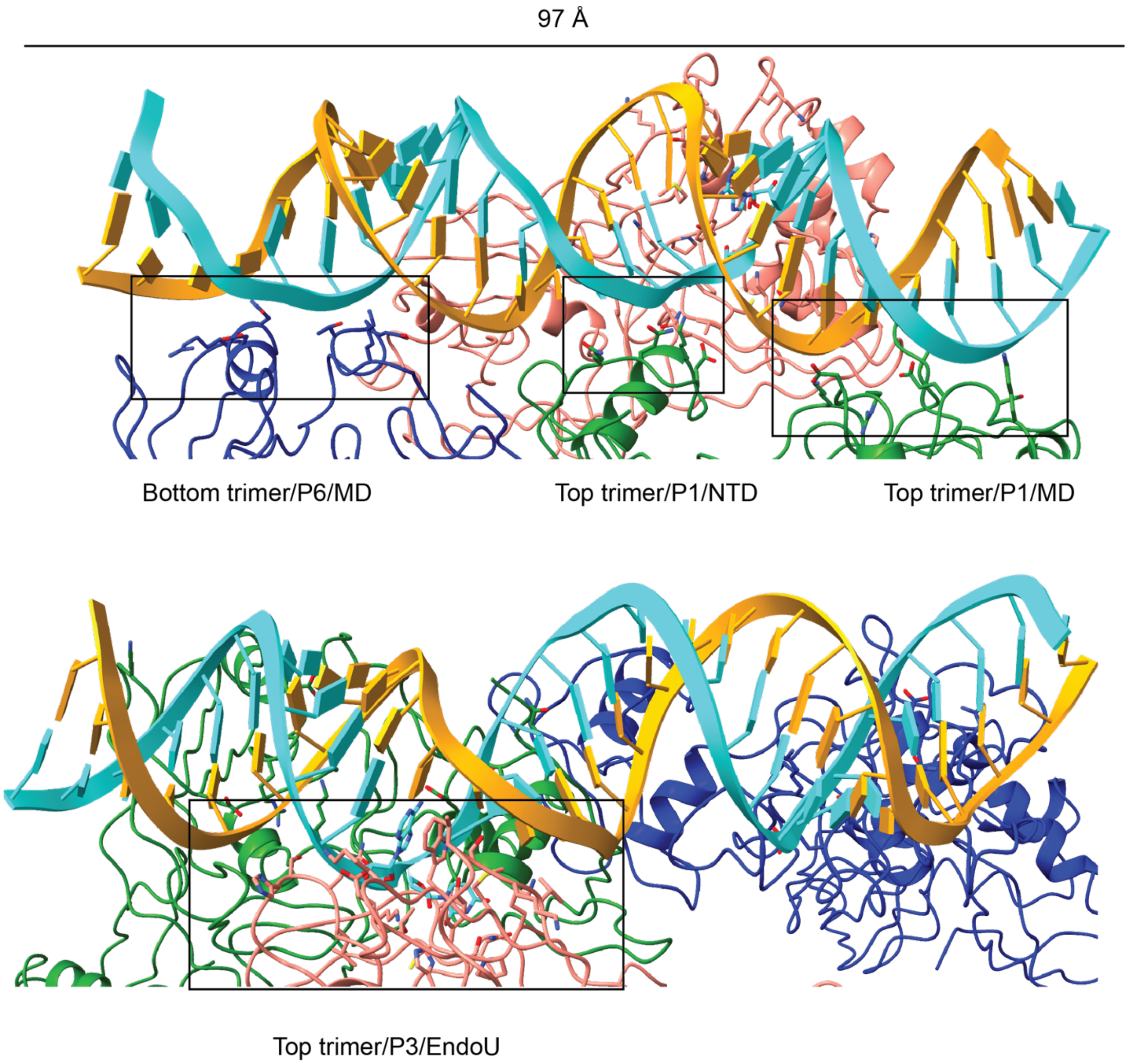


**Supplementary Figure 4. Nsp15/RNA interfaces.** Nsp15 interacts with dsRNA via 4 interfaces. The interfaces are denoted by black boxes, and the trimer position, protomer, and corresponding domain are listed. NTD=N-terminal domain, MD=middle domain, EndoU=catalytic C-terminal domain.

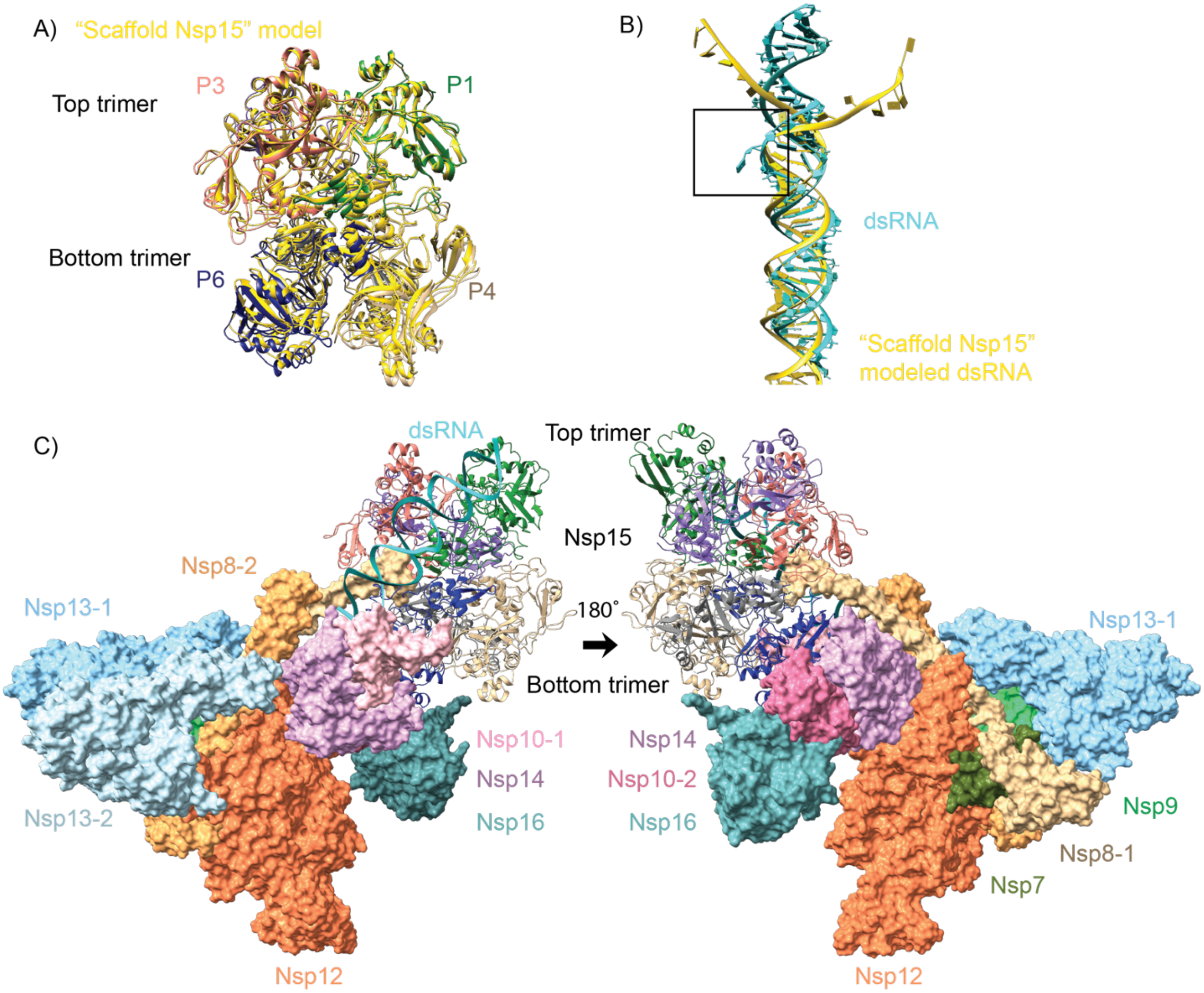


**Supplementary Figure 5. Scaffold model comparison with Nsp15/dsRNA structure.** A) The Nsp15 hexamer from our Arctica dataset Nsp15/dsRNA cryo-EM structure aligns well with the Nsp15 modeled as a scaffold in the RTC. (yellow). B) The dsRNA modeled into our Arctica dataset (cyan) aligns well with the dsRNA modeled into the RTC complex (yellow). Our experimental data show density for a uridine in an EndoU active site (black box). C) Extended view of the Nsp15 scaffold model (surface view[16] with our Nsp15/dsRNA structure (ribbon). For simplicity, only 1/6 of the scaffolded proteins are shown. Nsp12, the RNA-dependent RNA polymerase is shown in orange. Nsp8 (two copies, light tan and yellow-orange) and Nsp7 (olive) are polymerase accessory factors. The two copies of Nsp13, the helicase, are shown in light blue and sky blue. Nsp9, an RNA binding protein, is shown in light green. Nsp14 (light purple), a proofreading exonuclease, Nsp16 (teal), a methyltransferase, and Nsp10 (two copies, light pink and pink), their cofactor, round out the RTC scaffold model.

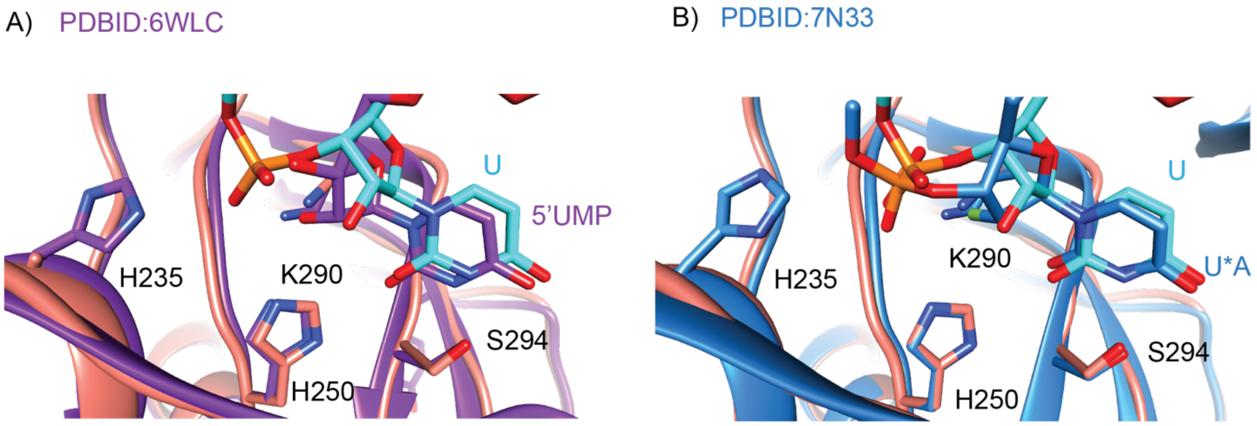


**Supplementary Figure 6. Active site comparisons.** A) Active site overlay of our cryo-EM structure (salmon/cyan) with an x-ray crystallography structure of Nsp15 with 5′-UMP (purple[17]). B) Active site overlay of our cryo-EM structure (salmon/cyan) with our previously determined pre-cleavage Nsp15 cryo-EM structure (blue[2]).

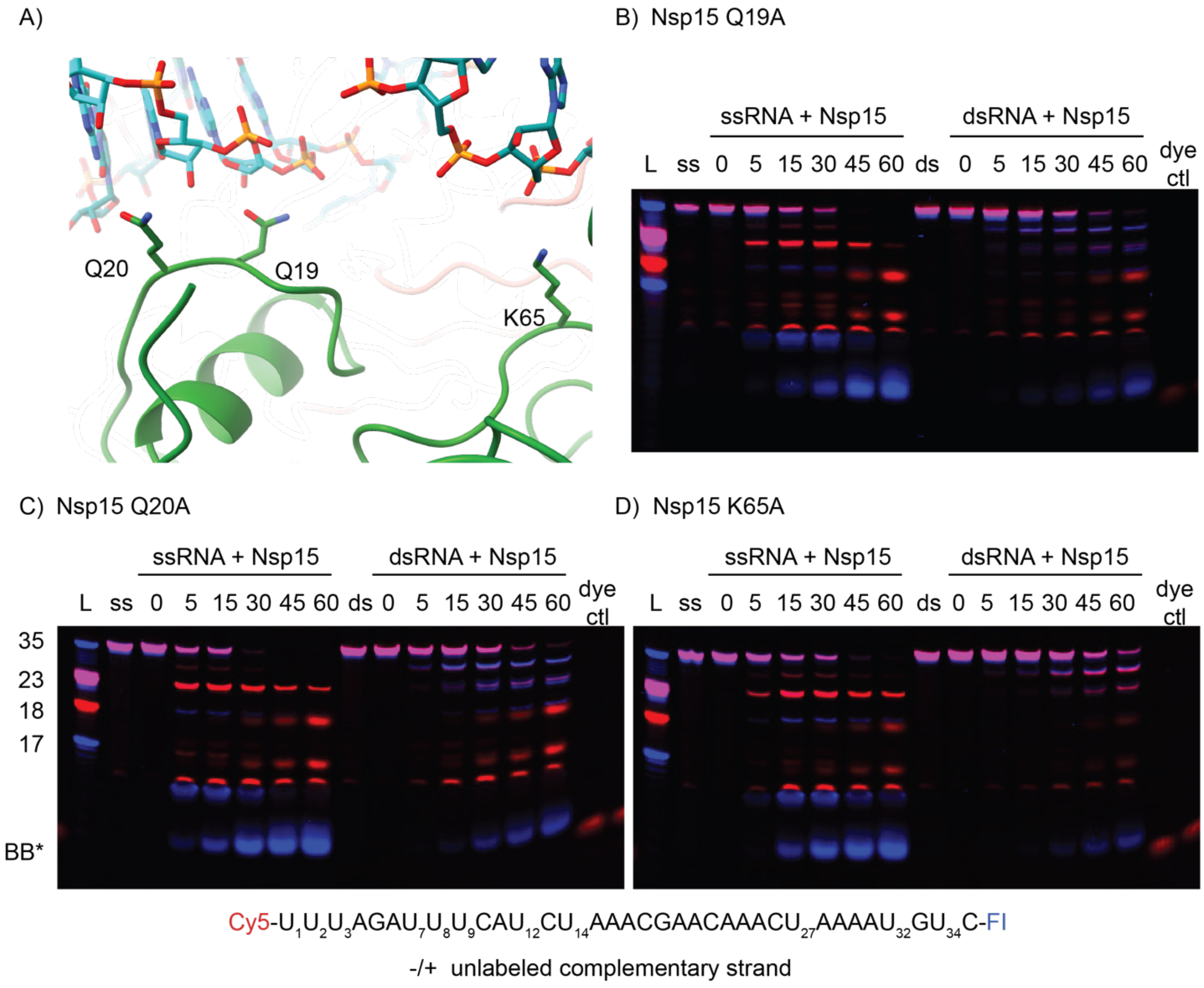


**Supplementary Figure 7. Representative, complete time course cleavage reaction gels comparing ssRNA vs. dsRNA for NTD/MD Nsp15 mutants. (A)** Zoomed in depiction of the NTD "platform" mutants. (**B-D**) 15% denaturing PAGE gels were used to resolve the nuclease assays. Ladder (L) nucleotide lengths are shown to the left of the gels. Bromophenol blue (BB*) runs ~8 nt in 15% denaturing gels.

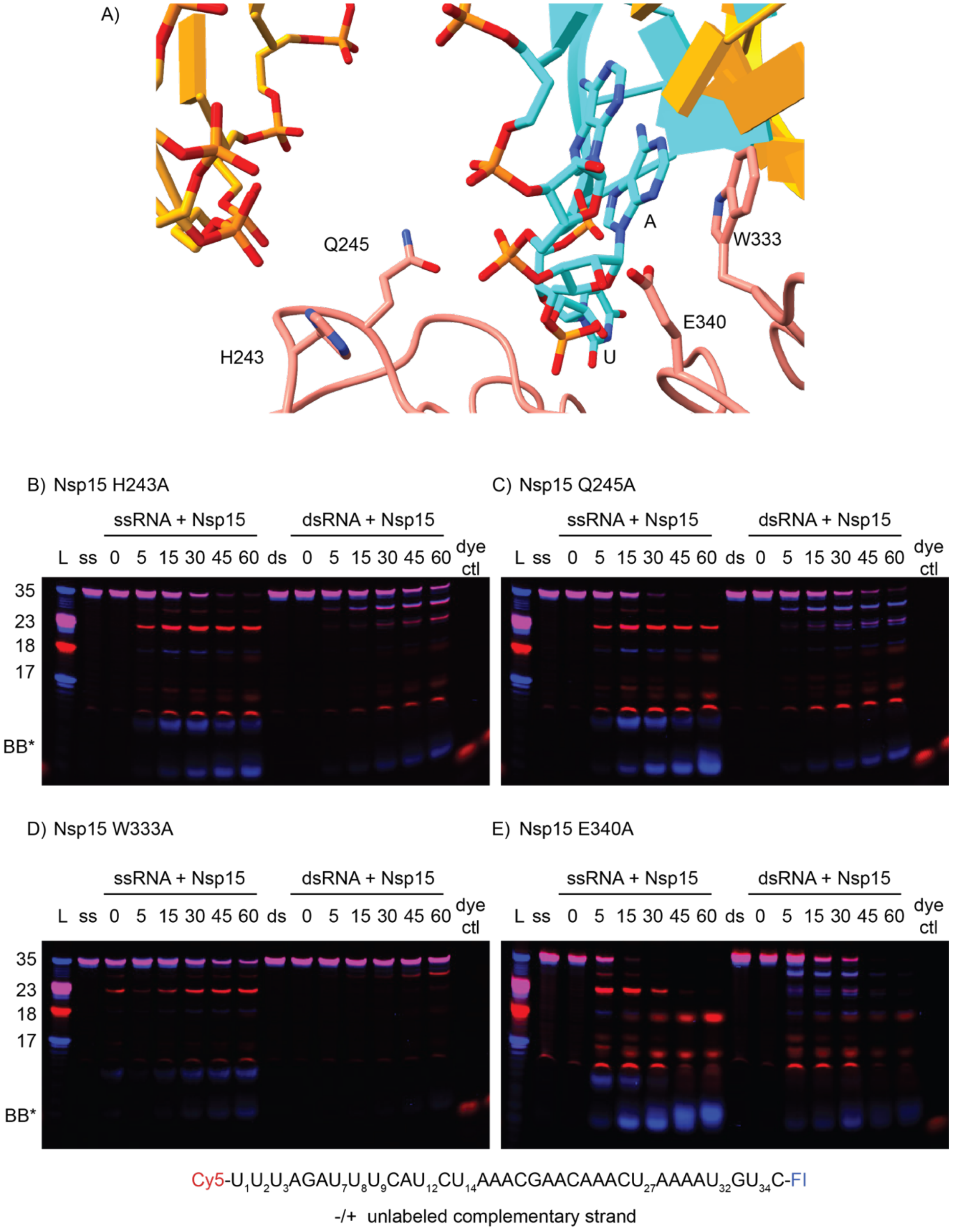


**Supplementary Figure 8. Representative, complete time course cleavage reaction gels comparing ssRNA vs. dsRNA for EndoU Nsp15 mutants. (A)** Zoomed in cartoon depiction of the active site, with mutated residues shown as sticks. **(B-D)** 15% denaturing PAGE gels were used to resolve the nuclease assays. Ladder (L) nucleotide lengths are shown to the left of the gels. Bromophenol blue (BB*) runs ~8 nt in 15% denaturing gels.

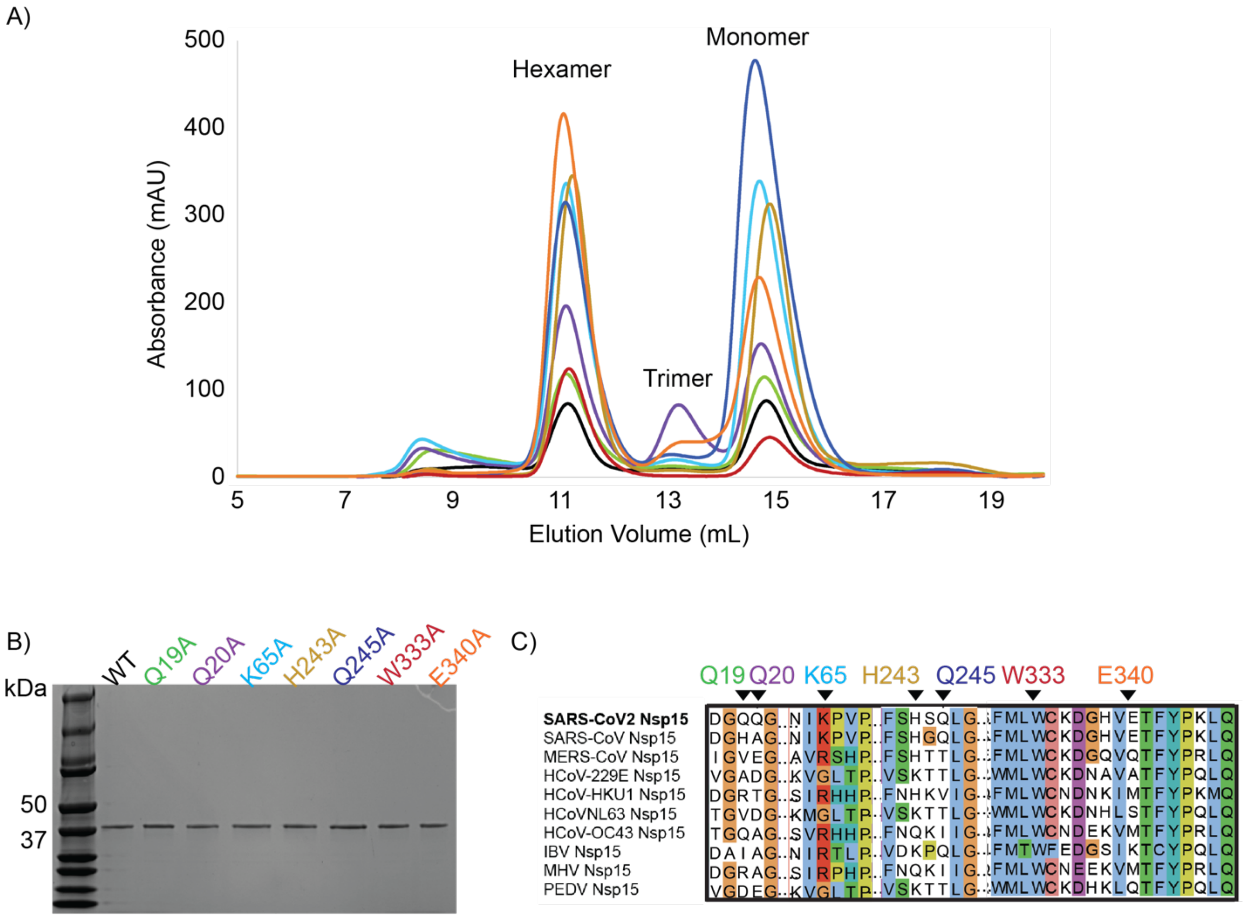


**Supplementary Figure 9. Biochemical characterization of Nsp15 mutants.** A) Size exclusion chromatography traces (S200 Superdex) for WT-Nsp15 and mutants. Two predominant peaks were seen for all proteins, corresponding to hexamer and monomer populations, although the proportion of each varied. Nsp15 Q20A also had a small peak corresponding to a trimer, which was inactive (data not shown). B) Summary SDS-PAGE gel for all Nsp15 constructs used in this study, showing pure protein. C) Sequence alignment for human coronaviruses and three major animal coronaviruses (infectious bronchitis virus, IBV; mouse hepatitis virus, MHV; porcine epidemic diarrhea virus, PEDV).

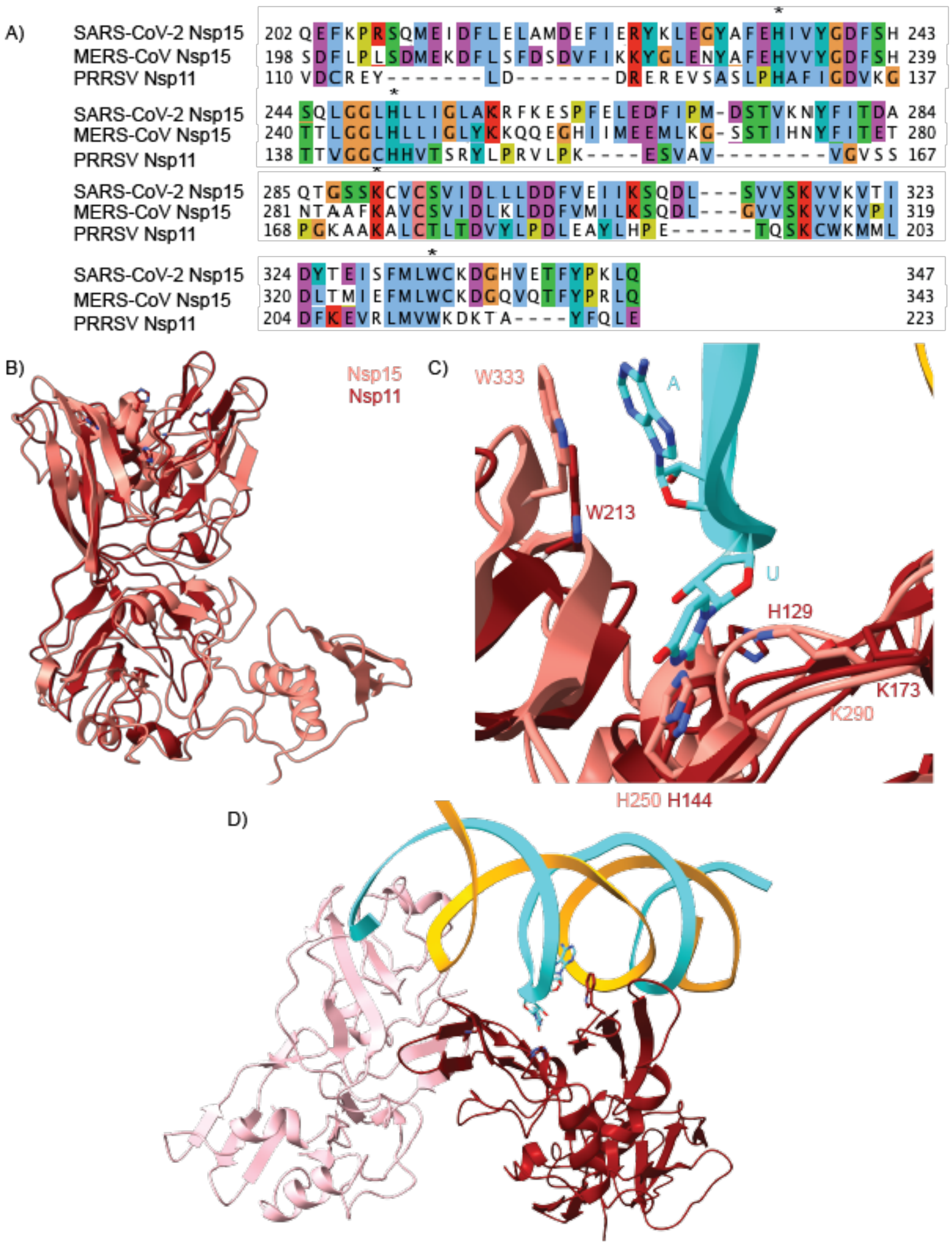


**Supplementary Figure 10. Conservation of dsRNA binding. A)** Sequence alignment of SARS-CoV-2 Nsp15, MERS-CoV Nsp15, and PRRSV (Porcine Reproductive and Respiratory Syndrome Virus) Nsp11. The conserved catalytic triad and π-stacking tryptophan are marked with asterisks. **B)** Superposition of protomers from SARS-CoV-2 Nsp15 (salmon, this study) and PRRSV Nsp11 (dark red, PDB: 5DA1[18]). The C-terminal EndoU domains superpose well around the active site, but otherwise the structures diverge. The overall RMSD is 7.2 Å. **C)** Zoom in of the active site overlay showing the catalytic triad and π-stacking tryptophan for Nsp11 and Nsp15 with dsRNA. **D)** The Nsp11 dimer with dsRNA modeled by superposition. The dsRNA is positioned to interact with the active site of one protomer and sits in a groove between the dimer.
